# Modified histone peptides uniquely tune the material properties of HP1α condensates

**DOI:** 10.1101/2024.02.07.579285

**Authors:** Priyasha Deshpande, Emily Prentice, Alfredo Vidal Ceballos, Patrizia Casaccia, Shana Elbaum-Garfinkle

## Abstract

Biomolecular condensates have emerged as a powerful new paradigm in cell biology with broad implications to human health and disease, particularly in the nucleus where phase separation is thought to underly elements of chromatin organization and regulation. Specifically, it has been recently reported that phase separation of heterochromatin protein 1alpha (HP1α) with DNA contributes to the formation of condensed chromatin states. HP1α localization to heterochromatic regions is mediated by its binding to specific repressive marks on the tail of histone H3, such as trimethylated lysine 9 on histone H3 (H3K9me3). However, whether epigenetic marks play an active role in modulating the material properties of HP1α and dictating emergent functions of its condensates, remains only partially understood. Here, we leverage a reductionist system, comprised of modified and unmodified histone H3 peptides, HP1α and DNA to examine the contribution of specific epigenetic marks to phase behavior of HP1α. We show that the presence of histone peptides bearing the repressive H3K9me3 is compatible with HP1α condensates, while peptides containing unmodified residues or bearing the transcriptional activation mark H3K4me3 are incompatible with HP1α phase separation. In addition, inspired by the decreased ratio of nuclear H3K9me3 to HP1α detected in cells exposed to uniaxial strain, using fluorescence microscopy and rheological approaches we demonstrate that H3K9me3 histone peptides modulate the dynamics and network properties of HP1α condensates in a concentration dependent manner. These data suggest that HP1α-DNA condensates are viscoelastic materials, whose properties may provide an explanation for the dynamic behavior of heterochromatin in cells in response to mechanostimulation.

**Statement of significance:** The organization of genomic information in eukaryotic cells necessitates compartmentalization into functional domains allowing for the expression of cell identity-specific genes, while repressing genes related to alternative fates. Heterochromatin hosts these transcriptionally silent regions of the genome - which ensure the stability of cell identity -and is characterized by repressive histone marks (H3K9m3) and other specialized proteins (HP1a), recently shown to phase separate with DNA. We show that HP1a forms condensates with DNA which persist in the presence of H3K9me3 peptides. The viscoelastic nature of these condensates depend on H3K9me3:HP1 ratios, which are modulated by mechanical strain in cells. Thus, phase separation may explain the dynamic behavior of heterochromatin in cells, in response to mechanostimulation.

## Introduction

Biomolecular phase separation has revolutionized the field of cell biology, with much interest directed towards understanding the role of condensate formation in genome organization within the eukaryotic nucleus (1–3). In mammalian cells, two meters of DNA containing precious genomic information are tightly packed within a membrane delimited organelle ranging between 5 to 10 microns in diameter. This size reduction is achieved by DNA condensation and wrapping of approximately 147 base pairs of DNA around positively charged octamers of histone proteins (dimers of H2A, H2B, H3 and H4) to form “core nucleosomes”, which represent the basic unit of chromatin (4), and repeat every 200 +/- 40 base pairs for the entire length of 6 billion bp of DNA (5). The intrinsically disordered N-terminal tails of core histone proteins emanate out of the nucleosome core particle and are subject to several post-translational modifications on specific amino acids (1). Linker histone H1 proteins bind to DNA outside the nucleosomes and have been shown to regulate the formation of “chromatosomes” (6), and to be further organized into higher order chromatin structures, with important functional and ultrastructural differences. Heterochromatin, characterized by an electrodense ultrastructure and condensed state, hosts transcriptionally silent/inactive regions of the genome, while euchromatin, which is characterized by an electrolucent appearance via electron microscopy and open/relaxed conformations of chromatin, hosts sites of active transcription (7). Heterochromatic compartments have been further classified into constitutive (containing permanently silent regions of the DNA such as centromeres and gene-poor DNA sequences), and facultative (containing cell and developmental-stage specific repressed genes). However, it has become increasingly clearly that, even constitutive heterochromatin needs to allow specific genomic regions to be locally accessed, either to allow for transcription (8) or to allow for DNA repair (9). The identification of specific sets of proteins that either deposit (<u>writers)</u>, or bind to (<u>readers</u>) or remove (<u>erasers</u>), specific post-translational modifications on lysine and arginine residues at the N terminal tails of H3 and H4 histones provided a potential mechanism for this regulation (10). At a molecular level, genomic sites of active transcription are characterized by high levels of trimethylation of lysine 4 (H3K4me3) at promoter regions, while sites of gene repression are characterized by high levels of H3K9me2/3, H3K27me3, H4K20me3 (11, 12). Since repression of alternative fates is crucial for somatic cell specialization, it is important to reconcile the fact that localized opening of specific genes must occur within an otherwise stable region of silenced chromatin. The initial discovery that the constitutive heterochromatin-enriched protein HP1α, has the ability to phase separate *in vivo* (13) and *in vitro* (14) provided the intriguing possibility that studying the behavior of condensates generated by heterochromatin-enriched proteins, would shed some light on the mechanism of eukaryotic genome organization and dynamic behavior. It was then shown that the mechanism of HP1α phase separation is highly conserved from yeasts (15) to humans (16) and that HP1α is capable of uniquely inducing DNA compaction (14, 17), thereby corroborating the importance of HP1α in contributing to heterochromatin formation (16). Several studies continued to characterize the biophysical requirement of HP1α phase separation, including the phosphorylation state of the protein (18), the importance of DNA length (17) and relation to nucleosomes (19). On the other hand, few studies addressed the ability of the intrinsically disordered histone tails to phase separate (20, 21) and led to the open question of whether and how histone marks on the tails of histone H3 may impact the phase separation behavior of HP1α, which we address in this study.

In addition, based on a growing literature supporting the importance of biophysical properties of chromatin, in modulating the response of cells to protect the genome from external mechanical stimuli (22) as well as recent evidence highlighting novel functions of HP1α in regulating chromosomal mechanics and stiffness (23, 24) we asked whether the repressive H3K9me3 mark fine tunes the viscoelastic material properties of heterochromatin condensates formed by HP1α and DNA. Here, we specifically leverage a reductionist approach to directly examine the interplay of the heterochromatic mark H3K9me3 with its reader protein HP1α in the context of phase separation and characterize the material properties of HP1α condensates.

## Materials and Methods

### Materials

Recombinantly expressed and purified HP1α (MW = 22kDa) was purchased (lyophilized against PBS buffer) from Biomatik Corporation (www.biomatik.com) (Ontario, Canada). Unmodified and methylated histone H3 peptides (1-22) (MW =2354) were purchased in lyophilized form from Bachem Americas,Inc (Torrance, CA) (<u>Bachem.com</u>) and used as received. Chemicals including Alexa-488 C5-Maleimide, Dylight-594 NHS ester, Nucleic acid dimer sampler kit (YOYO-3 iodide, POPO-1 iodide), dsDNA (NoLimits 2.5kbp, 4kbp, 10kbp) as well as dialysis cassettes (Slide-A-Lyzer G2 3kda cutoff) were obtained from Thermo Fisher Scientific (Waltham, MA). Potassium chloride, Pluronics F127 was obtained from Sigma Aldrich (St. Louis, MO). HEPES ((4-(2-hydroxyethyl)-1-piperazineethanesulfonic acid)) was obtained from Fisher Scientific (Pittsburg, PA). Peptides were reconstituted at 25 mg ml^-1^ in nuclease-free ultra pure water as per the manufacturer’s instructions and stored as aliquots at -20°.HP1α was reconstituted (0.5 mg ml^-1^) in nuclease-free ultra pure water as per the manufacturer’s instructions and stored (with 10% glycerol) as flash-frozen 200 ul aliquots in -80°C.

#### Protein labeling

Reconstituted HP1α was buffer exchanged into 70mM KCl, 20mM HEPES, 1mM TCEP, pH 7.4 buffer before setting up the labeling reaction. HP1α was fluorescently labeled with Invitrogen alexa-488 C5-Maleimide per the manufacturers instructions. Briefly, ∼150-200uM of protein was mixed with 10-fold molar excess solution of dye and the reaction was incubated for 30 minutes at room temperature, protected from light. The reaction was quenched by adding 10mM DTT and labeled protein was separated from dye dialysis using 3kDa cutoff dialysis cassettes. Protein was concentrated using a 3kDa cutoff Amicon ultra-centrifuge filter and was flash frozen (with 10% glycerol) and stored at -80°C. Concentration of labeled protein was determined using absorbance at 488 nm.

#### Peptide labeling

Histone H3 peptides (unmodified, K9/K4me3-modified) (N = 22) were labeled with Dylight594 NHS ester as per the protocol provided by the manufacturer. Briefly, 200uL of 2mg/ml solution of peptide was mixed with 10-fold molar excess solution of dye and incubated at room temperature for 1-2 hours. The labeled peptide was separated from free dye by dialysis using 1kDa Molecular Weight Cut Off Pur-A-Lyzer midi dialysis cassette (Sigma Aldrich). To make sure all the dye is removed, the peptide solution was washed with 70mM KCl buffer in a 3kDa Amicon filter till the flowthrough absorbance (594nM) reached 0. Labeling was confirmed using Fluorescence Correlation Spectroscopy measurements to determine the diffusion coefficient of labeled peptide in comparison to free dye and by LC-MS. Degree of labeling was calculated using absorbance at 594 nm and 205 nm (peptide backbone absorbance).

#### DNA labeling

dsDNA (2.5kbp, 4kbp, 10kbp) was labeled with YOYO-3 (612/631 nm) or POPO-1 (434/456 nm) dyes before every experiment. Briefly, 1mM stock of the dye (in DMSO) was diluted 1:500 (v/v) into 70mM KCl, 20mM HEPES, 1mM DTT, pH 7.5 buffer. Depending upon the volume of DNA required for the experiment, the DNA was added to the diluted dye solution and incubated for 1-2 mins. Phase separation experiments were performed by using ∼3%labeled DNA mixed with unlabeled DNA to maintain the final concentration of DNA constant. Care was taken to maintain at least a 5:1 DNA:dye molecules ratio by adjusting the dye substock concentrations depending upon the experiment.

#### Coacervate preparation

Prior to coacervate preparation, HP1α was filtered using 0.22um filter and buffer exchanged 70mM KCl, 20mM HEPES, 1mM DTT, pH 7.4. The protein was concentrated using a 3kDa cutoff Amicon ultra centrifugal filter unit until a desired concentration (∼80-100uM) was achieved. The microscope glass slide used for reaction incubation was cleaned using 70% ethanol and dried, followed by treatment with Pluronics F-127 treatment for 1 hour. The slide was then washed five to six times using deionized water.

Coacervate samples were prepared by mixing required amounts of protein and dsDNA (and peptides) to achieve a final concentration of 20uM HP1α and desired concentrations of DNA and peptide. Final volume of the reaction mixture was kept constant at 7ul. The reaction temperature was maintained at 23°C (CherryTemp, CheeryBiotech).

#### DIC and Confocal imaging

DIC images were acquired using a wide-field Axio Observer 7 Inverted Microscope (Zeiss) with a ×40 numerical aperture (NA) Plan-Apochromat air objective. For confocal imaging, samples were imaged using a Marianas Spinning Disc confocal microscope (Intelligent Imaging Innovations) which consisted a spinning disc confocal head (CSU-X1; Yokagawa) and ×63/1.4-NA Plan-Apochromat (oil immersion) objective. Alexa 488 was excited using a 488-nm line from a solid state laser (LaserStack) while Dylight-594 was excited with a 561-nm line and POPO-1 was excited at 405 nm line from the same laser. Emissions were collected using a 440/521/607/700nm quad emission dichroic and 525/530 nm emission filter. The 561-nm line was used for imaging 200-nm carboxy beads used in microrheology experiments.Image acquisition was done using a Prime sCMOS camera (Photometrics), controlled by Slidebook 6 (Intelligent Imaging Innovations). ImageJ software was used to format, crop and analyze images.

#### FRAP

Fluorescence Recovery After Photobleaching experiments were performed for Alexa-488 labeled HP1α as well as YOYO-3 labeled dsDNA. Briefly, a small amount (∼1%) of labeled protein/DNA is mixed with unlabeled protein/DNA before inducing coacervation. Final concentration of protein was 20uM, DNA concentration varied depending upon the length used from 40nM (2.5kbp) to 9nM (10kbp). A 0.5um-1um sized area is selectively photobleached and the recovery of fluorescence is tracked over time. FRAP images were acquired using spinning disc confocal microscope (Intelligent Imaging Innovations). Images were acquired at 200ms timestep and 50-100ms exposure time and recovery was monitored for 200-300 points. Alexa 488 was excited using the 488nm line from solid state laser (LaserStack) while YOYO-3 was excited using the 561nm line. Emissions were collected using 440-/521-/607-/700-nm quad emission dichroic and 525-/30-nm emission filter. The recovery of fluorescence was plotted as a function of time and fitted using exponential function of the form -

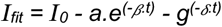

ImageJ was used to process the images and the recovery of fluorescence was normalized using MATLAB to obtain the mobile fraction (I_0_) and half-time of recovery [*ln*(2)/*β*)].

#### Microrheology

200nm-carboxylated modified fluorescent beads (Invitrogen) were embedded in HP1α-DNA or HP1a-DNA-H3K9me3 droplets and their motion was tracked as a function of time with 200ms interval for 1000 frames with 10-50ms exposure. Bead diffusion was tracked using Marianas spinning disc confocal microscope with a x63/1.4NA Plan-Achromat oil objective. Temperature of the sample was maintained at 23°C ± 2°C using a microfluidic temperature stage (CherryTemp, CherryBiotech). Care was taken to select beads away from coarvate boundaries to avoid boundary effects. Particle-tracking code to locate and trace particle trajectories in two-dimensions (XY) was adapted from MATLAB Multiple Particle Tracking Code from The MATLAB particle tracking code repository(https://www.pnas.org/doi/full/10.1073/pnas.1504822112, https://doi.org/10.1038/s41467-020-18224-y). Bead dynamics was analyzed using custom MATLAB software. Mean Square Displacement of the beads was calculated from time and ensemble averages for all trajectories :

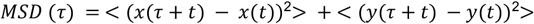

The relationship between the MSD and the lag time (*τ*) follows a power law, the diffusive exponent (α) was determined as the slope of the log-log plot.

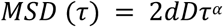

Where d is the number of dimensions (here 2) and D is the diffusion coefficient. If the value of diffusive exponent (*α*) is not equal to 1, the diffusion coefficient cannot be used to extract the value of viscosity. The values of α less that 1 indicate that the material is not a pure, viscous liquid but is viscoelastic in nature.

#### Fluorescence Correlation Spectroscopy Measurements-

Histone H3 vs. H3K9me3 peptide binding to HP1α was quantified using Fluorescence Correlation Spectroscopy measurements to measure translational diffusion of fluorescently labeled histone peptides with increasing concentrations of HP1α protein at 23°C.

FCS binding measurements were collected with an inverted Leica TCS SP8 STED 3X using a 63x water immersion objective. The fluorophores were excited at 561 nm with a white light laser and detected between 571 – 581 nm using a HyD detector. Data acquisition and analysis were performed using SymPhoTime (PicoQuant, Germany) software, with each measurement being an average of three 300 s traces to obtain the correlation curves G(*τ*). The average of these correlation curves were fit to a single-component triplet model using the following equation:

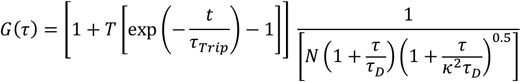

where G(*τ*) is the autocorrelation function as a function of time, *τ*. N is the average number of molecules in the focal volume. *τ*_*D*_ is the average amount diffusion time of fluorescent molecules through the focal volume. 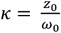 represents the ratio of the axial *z*_0_ and radial dimensions *ω*_0_ of the Gaussian excitation volume. T is the dark (triplet) fraction of molecules and *τ*_*Trip*_ is the lifetime of the dark (triplet) state. The excitation volume was determined using sulforhodamine B dye in water, with a diffusion coefficient of 4.3 × 10^−6^*cm*^2^*s*^2^ at 25 °C. The measured *τ*_*D*_ of the fluorescent molecules at increasing protein concentration was plotted and fit using Prism 9 using the one site - total binding model:

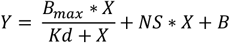

where *B*_*max*_ is the maximum specific binding, Kd is the equilibrium dissociation constant in nM, and NS is the slope of non-specific binding.

#### Cell Tension Experiments

To test the effects of uniaxial strain on H3K9me3 and HP1a levels in cell nuclei, BV2 microglia cells were seeded at a density of 15,000 cells/cm^2^ on commercially available tissue culture plates (Flexcell International, Uniflex) with elastomer bottoms pre-treated with laminin for use in the Flexcell tension system (Flexcell International, T-5000). Immediately following completion of the tension regimen (12% elongation, 0.1Hz, sine wave for 30 min), immunocytochemistry was performed as previously described (25). Click or tap here to enter text.Both strained culture plates and unstrained control plates were incubated with primary antibodies to detect H3K9me3 (Diagenode, C15200146) and HP1a (Abcam, ab109028), followed by incubation with secondary antibodies tagged with Alexa fluorophores (Invitrogen, A11034; Invitrogen, A21422). Samples were mounted with media containing DAPI (Southern Biotech, Fluoromount with DAPI-G) and visualized using a Leica Scanning Confocal Microscope (LSM 800). All images were acquired with constant and defined laser parameters. Quantitative analysis of medial optical slices of cell nuclei was performed using FIJI (NIH), and all statistical analysis was performed in Prism (Graphpad).

## Results

### HP1α forms dynamic, viscoelastic condensates with DNA sequences of different length

Previous work has demonstrated that even in the absence of the H3K9me3 mark, HP1α can condense thousands of base pair long stretches of DNA via an underlying phase separation mechanism (17). In the resulting phases, HP1α remains highly dynamic and is rapidly exchanged between condensates upon fusion while DNA remains compacted and confined to specific territories. Therefore, to examine whether the repressive H3K9me3 mark impacts the phase behavior of HP1α-DNA, we first characterized phase separation of unmodified, human HP1α with DNA using three different, fixed lengths of DNA (N = 2500 bp, 4000 bp and 10000 bp). The selected sequences do not code for any genes. We observe that HP1α undergoes phase separation at near physiological concentrations (12.5μM-20μM tested here) with all three DNA lengths at nanomolar concentrations (70mM KCl, 20mM HEPES, 1mM DTT, pH 7.5) (Figure 1A). Phase separation is induced at lower concentrations of DNA with an increase in the length of the DNA monomer, ranging from 30nM for 2500 bp to 9nM for 10000 bp. HP1α, in the absence of DNA, remains soluble at the same concentration (Figure 1A). While spherical droplets appeared instantaneously for 2.5kbp and 4kbp DNA (Figure 1B), in the case of 10kbp DNA, assemblies with an asymmetric morphology were initially observed before transitioning to spherical droplets within approximately 1 hour (SI Movie 1) (Figure 1C). FRAP measurements with ∼1% Alexa48-labeled HP1α revealed that HP1α molecules remain dynamic within HP1α-DNA condensates for all three DNA lengths, with ∼98% recovery and a t1/2 of ∼5 Sec (Figure 1D, Supplemental Figure 1). Similar FRAP measurements with ∼0.5% YOYO3-labeled nuclei acid, indicated that DNA molecules remain significantly less dynamic (Figure 1D, Supplemental Figure 1). Since FRAP only reports on the specific dynamics of the fluorescent molecule probed, we chose to further characterize the emergent material properties of the condensates using microrheology, a technique in which the motion of fluorescent tracer particles embedded within a condensate, is tracked over time (Figure 1E). The viscoelastic properties of the condensates can be extracted from analysis of the Mean Squared Displacement (MSD) of the particle trajectories (26). MSD varies linearly with time in the case of pure viscous fluids lacking any elastic component, with the slope of the fitted MSD versus time plot, known as the diffusive exponent (α) is equal to 1. The value of the diffusive exponent <1 indicates sub-diffusive motion and is associated with the presence of an elastic component. For rheological measurements performed with 20 μM HP1α and 40nM 2.5kbp DNA, we measured the diffusive exponent α to be 0.85±0.06, consistent with sub-diffusive behavior in a viscoelastic material. The average α value decreased with increasing DNA length, ranging from 0.72±0.02 for those with 4kbp of DNA to 0.58±0.05 for those with 10 kbp DNA (Figure 1F) and suggest an increase in the elastic component of condensates as a function of DNA length. Thus, even though HP1α and DNA remain dynamic within the condensates, the resulting phases are not pure liquids but are viscoelastic in nature.

**Figure 1.**
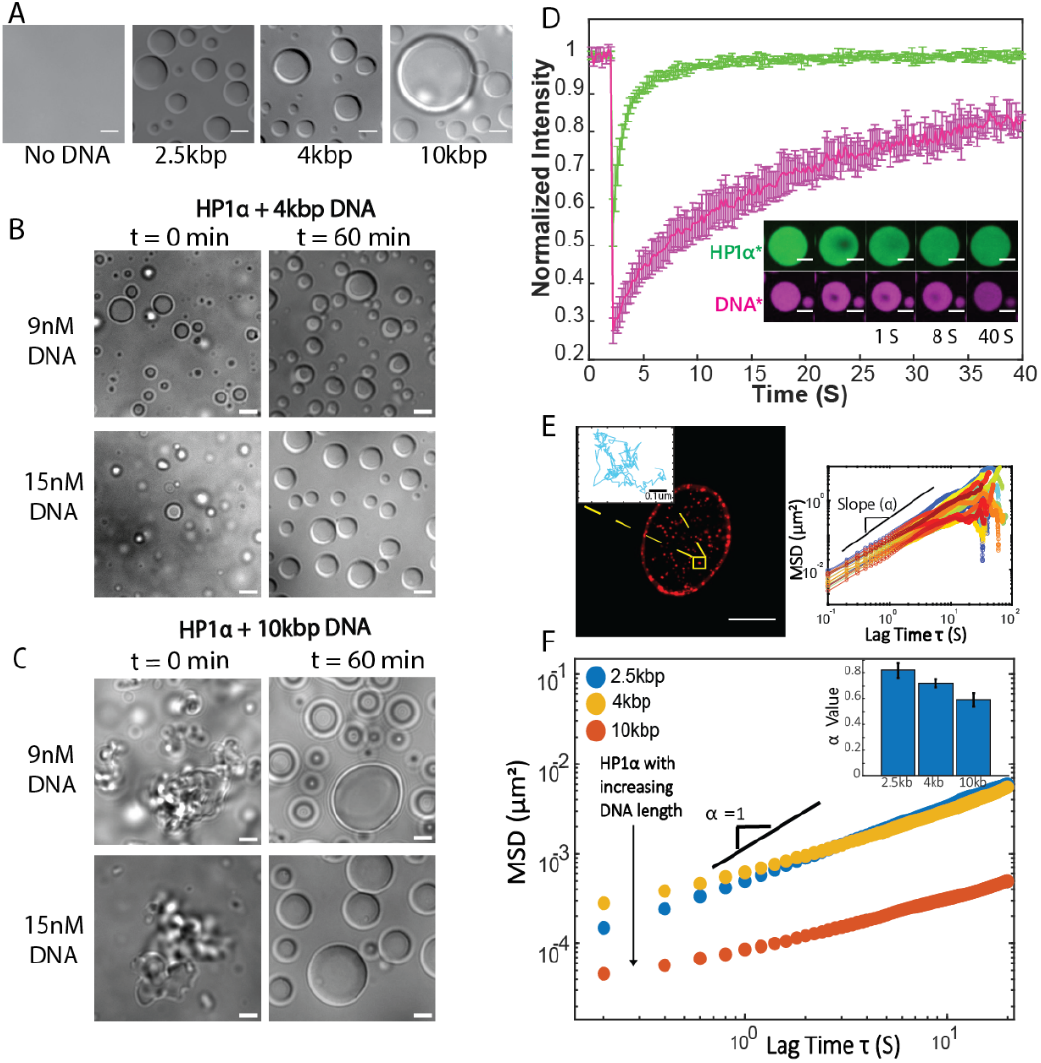
HP1α forms dynamic, viscoelastic condensates with DNA. **A)** DIC images (magnification = 40x) of HP1α-DNA mixtures at 20μM HP1α and 40nM 2.5kbp, 20nM 4kbp and 9nM10kbp DNA (70 mMKCl, 20 mM HEPES, 1 mM DTT, pH 7.5, Scale =5 μm)at 1hr incubation time. **B)** DIC images (magnification = 40x) of 20μM HP1α with 9nM (top) and 15nM (bottom) 4kbp DNA taken at incubation timepoints t=0min and t=60min. At both the timepoints spherical condensates can be seen. Scale bar = 5 μm. **C)** DIC images (magnification = 40x) of 20μM HP1α with 9nM (top) and 15nM (bottom) 10kbp DNA taken at incubation timepoints t=0min and t=60min. The images show changes in the morphology (from asymmetric to spherical) of condensates over a period of 1hour. Scale bar = 5 μm. **D)** Fluorescence Recovery After Photobleaching (FRAP) comparison for Alexa-488 20 μM HP1α(Green) and 40nM YOYO-3 labeled 2.5kbp DNA (Magenta). Scale = 5μm. **E)** Schematic showing distribution of beads in HP1a-DNA droplets and corresponding mean squared displacement (MSD) vs lag time plot for individual bead (200nm,Red FluoSpheres, Invitrogen). **F)** Mean Squared Displacements (MSD)vs. lag time for 20 μM HP1α condensates with 40nM 2.5kbp (blue), 20nM4kbp (yellow) and9nM10kbp (red) DNA. Inset, diffusive exponent (α) as a function of DNA length.

### Histone peptides with post-translational modifications of lysine residues associated with heterochromatic repression, such as H3K9me3, preserve the ability of HP1α to form condensates with DNA

The nuclear organization of the genome in mammalian cells is characterized by the enrichment of specific histone marks related to distinct transcriptional states, with repressive H3K9me3 enriched in heterochromatic regions and H3K4me3 enriched at promoter regions of transcriptionally active genes. HP1α, via its chromodomain, recognizes and binds specifically to the H3K9me3 mark on the N-terminal tail of histone H3, but does not bind to unmodified H3 or other trimethyl marks such as H3K4me3 (27). Consistently, immunohistochemical analysis of BV2 cells, an immortalized microglial murine line (Figure 2A) revealed a characteristic distribution of the repressive H3K9me3 mark at the nuclear periphery and in chromocenters, characterized by highly condensed DAPI+ DNA and co-localization with HP1α (Figure 2B), while the transcriptionally active H3K4me3 mark was entirely excluded from chromocenter immunoreactivity (data not shown). Since genome organization implies the coexistence of euchromatic and heterochromatic regions, it is conceivable that the ability of HP1α to form condensates may be affected by the presence of both transcriptionally repressive H3K9me3 and transcriptionally active H3K4me3 marks. For this reason, we sought to characterize the relative contribution of histone peptides containing these modifications or unmodified, in regulating phase separation of HP1α with DNA. To determine whether distinct post-translational histone modifications could directly influence the phase separation properties of HP1α-DNA condensates, we examined the phase behavior of unphosphorylated human HP1α and DNA in the presence of modified and unmodified segments of the human N-terminal histone H3 tail (residues 1-22). We used human HP1 alpha proteins, whose purity and molecular weight were confirmed using gel electrophoresis and Coomassie staining (Supplemental Figure S2A), and histone peptides whose molecular weight was confirmed using MALDI (Supplemental Figure S2B). Using Fluorescence Correlation Spectroscopy, we also showed the greater binding affinity of HP1a to histone peptides containing the repressive H3K9me3 mark, compared to unmodified histone H3 (Supplemental Figure S2C). Under identical experimental conditions, in terms of salt and protein concentration, the presence of H3K9me3 tails was compatible with the formation of phase separated condensates (Figure 2C and SI Movie 2-3). Unmodified H3 tails, in contrast, or tails containing marks of transcriptionally active genes, such as H3K4me3 abrogated HP1α-DNA phase separation for all three lengths of DNA tested, leading to the formation of aggregates (Figure 2C). In addition, similar to the results in BV2 cells, confocal imaging of fluorescently labeled components confirmed that HP1α, histone H3K9me3 peptide and DNA colocalized in the tri-component system *in vitro* (Figure 2D). The aggregated assemblies obtained with unmodified H3 tails, in contrast, were composed of histone peptides and DNA and lacked HP1α (Figure 2E, SI Movie 4) with no detectable FRAP recovery of DNA (Supplemental Figure S3). Notably, in the absence of DNA both modified and unmodified histone peptides did not promote HP1α phase separation, while in the absence of HP1α, peptides assembled into aggregates at all the concentrations tested (Supplemental Figure S4, SI Movie 4, SI movie 5).

**Figure 2.**
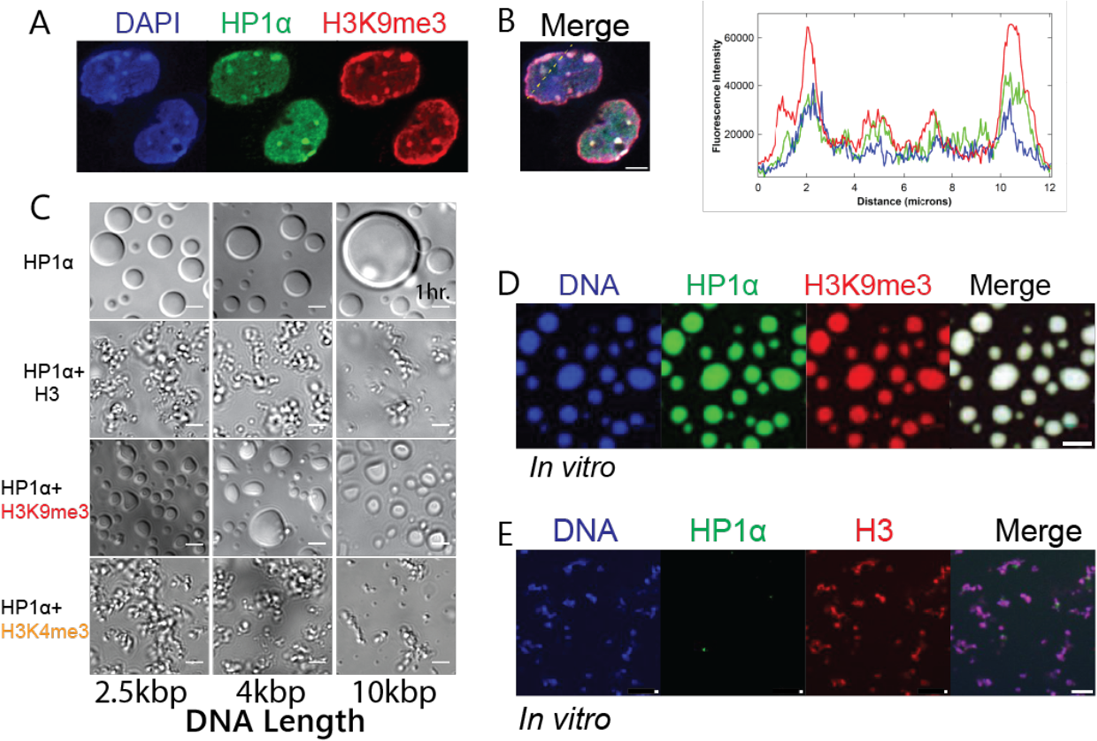
Histone peptides containing the heterochromatic H3K9me3 mark are compatible with the formation of HP1α condensates with DNA, while unmodified or H3K4me3 containing peptides are not. **A)** Confocal image of murine BV2 microglia nuclei stained with antibodies specific for the heterochromatic components HP1α (green) and H3K9me3(red). DAPI (blue) was used to label DNA. Scale = 5μm. **B)** This representative line plot shows the relative immunoreactive profile across eukaryotic nuclei revealing the colocalization of HP1α (green), H3K9me3 (red) and DNA (blue). **C)** DIC images of HP1α-DNA mixtures at 20μM HP1α and 40nM 2.5kbp, 20nM 4kbp and 9nM10kbp DNA without (top row) and with 42μM unmodified and modified histone peptides (70mMKCl, 20mM HEPES, 1mM DTT, pH 7.5, Scale =5μm) at 1hr incubation time. **D)** Confocal images (Magnification = 63X) of in vitro reconstituted condensates showing colocalization of HP1α(20μM)(Green), DNA(2.5kbp 40nM)(Blue) and H3K9me3 peptide(42μM)(Red). Scale = 5μm. **E)** Confocal fluorescence images of in vitro assemblies of HP1α(20μM)(Green), DNA(2.5kbp 40nM)(Blue) and unmodified H3 peptide(42 μM)(Red) (1-22). Scale = 5μm.

### Dynamic changes of heterochromatic mark distribution in BV2 cells in response to mechanical strain

It is important to highlight the fact that heterochromatin, despite its condensed structure in nuclei, is also a dynamic compartment, with the ability to respond to signals provided to biological systems. From a functional perspective, the localization of the heterochromatic H3K9me3 mark at the nuclear periphery, underneath the nuclear envelope (28–31), has been proposed to have a protective role for the genome, by defending the genomic integrity (24, 32). However, as cells are challenged by external physical forces, such as mechanical strain, it is important that heterochromatic regions may dynamically adjust in terms of flexibility, as a rigid structure would result in brittle nuclei and shattered DNA in response to mechanical stressors (32). To define the effect of mechanical strain on heterochromatic components we subjected BV2 cells to uniaxial strain for 30 minutes and evaluated the consequences on the nuclear distribution of H3K9me3 and HP1α in the cell nuclei, using immunocytochemistry with specific antibodies and used DAPI as DNA counterstain to identify chromocenters. Consistent with the localization of heterochromatin in differentiated cells, unstrained BV2 cells were characterized by the presence of the histone H3K9me3 mark at the nuclear periphery and in DAPI+ chromocenters (Figure 3A), which co-localized with the immunoreactivity for HP1α (Figure 3B). In response to uniaxial strain, however, the immunoreactivity for H3K9me3 decreased (Figure 3C-D) both at the nuclear periphery and at internal chromocenters (Figure 3E), while HP1α nuclear immunoreactivity remained unchanged (Figure 3F). These data suggest a strategy by which cells modulate the relative HP1α:H3K9me3 ratio in response to mechanical forces.

**Figure 3:**
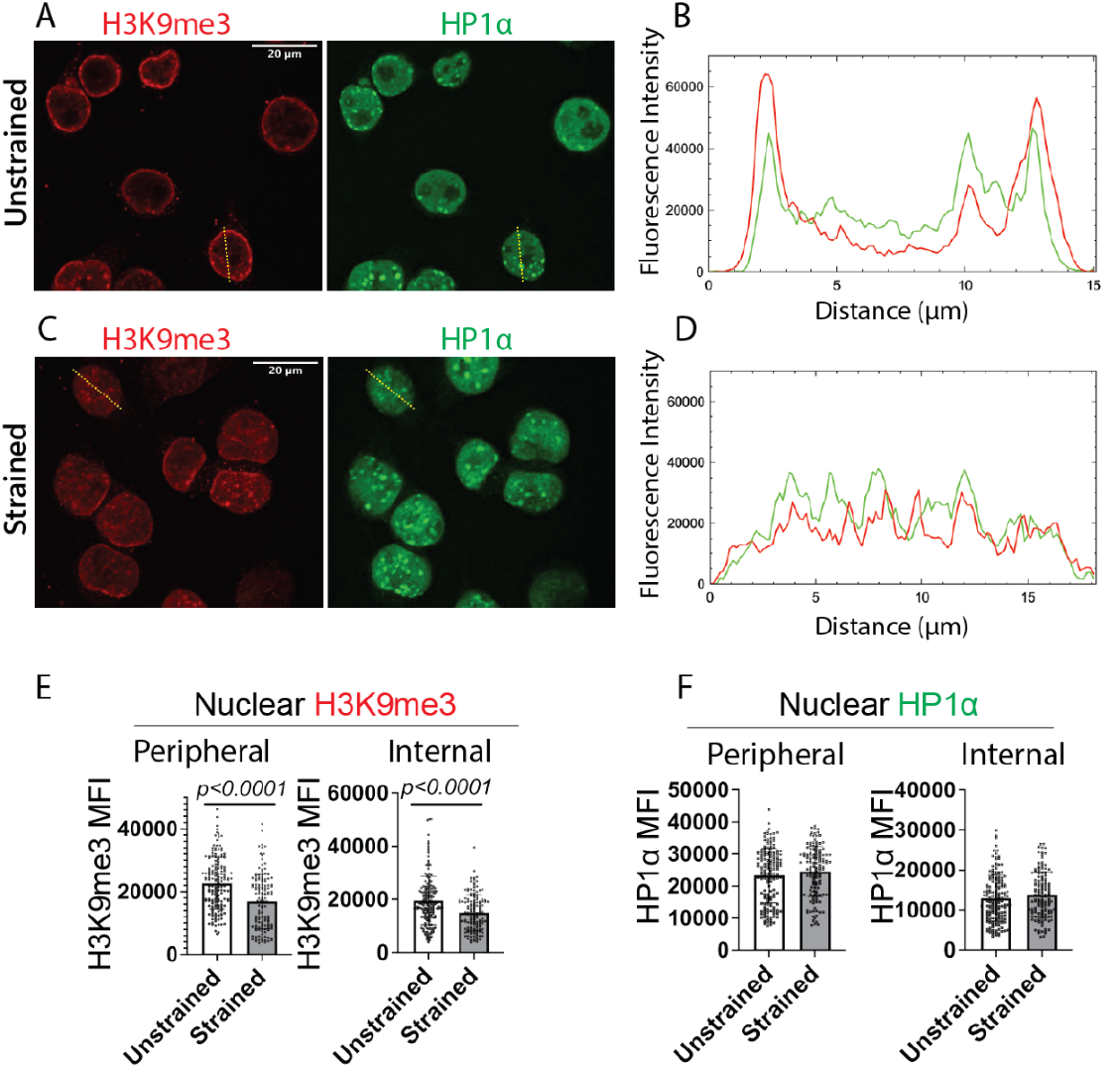
Uniaxial strain of mammalian microglial cells results in decreased H3K9me3 concentrations at nuclear periphery and chromocenters. **A)** Confocal images of unstrained BV2 cell nuclei fixed and stained with antibodies for H3K9me3 (red) and HP1α(green). Scale = 20μm. **B)** Line profile across a selected cell nucleus of an unstrained BV2 cell indicating mean fluorescence intensity of HP1α (green) and H3K9me3 (red) indicating strong colocalization between the two components at the nuclear periphery. **C)** Confocal images of BV2 cell nuclei exposed to uniaxial tensile strain for 30 minutes, then fixed and stained with antibodies for H3K9me3 (red) and HP1α(green). Scale = 20μm. **D)** Representative line profile of HP1α (green) and H3K9me3 (red) immunoreactivity across a selected cell nucleus of a strained BV2 cell revealing decreased level of H3K9me3 fluorescence intensity. **E, F)** Graphs quantifying the relative mean fluorescence intensity (MFI) of H3K9me3 (E) and HPIα (F) at the nuclear periphery and in internal chromocenters in the nuclei of strained (n=134 cells) and unstrained (n=166 cells) from four independent biological replicates. Each dot represents an individual cell. Two-way Student’s t-test.

### H3K9me3 mark tunes HP1a condensate properties and dynamics

The decreasing levels of H3K9me3 in BV2 cells, in response to mechanical strain, while HP1α concentrations remained constant, further prompted us to evaluate the effect of the relative levels of H3K9me3 onto the phase separation properties of HP1α /DNA condensates. We therefore tested the effect of a range of H3K9me3 peptide concentrations on the ability of HP1α to phase separate and on the material properties of these condensates We noted that concentrations of the H3K9me3 tail ranging from 8μM-42μM did not disrupt the condensate morphology (Figure 4A). We then asked whether the same range of concentrations of the H3K9me3 peptide influenced the internal dynamics and network properties of HP1α condensates. The use of FRAP allowed us to identify an alteration of the internal dynamics of HP1α within condensates which was dependent on the relative concentration of the H3K9me3 histone peptide (Figure 4B). At the highest concentration of H3K9me3 peptide tested (42μM), the mobile fraction of HP1α was ∼70% compared to ∼100% at lower concentration of the peptide (10μM). For condensates comprising HP1α /H3K9me3/ 2.5kbp DNA, we observed a significant decrease in the value of diffusive exponent (***α***) - from ∼0.85±0.06 to ∼0.3±0.03 for both concentrations tested (Figure 4C), which is close to the noisefloor or resolution threshold of our measurment. The decrease in ***α*** observed for the H3K9me3 mark with 2.5kb DNA was more pronounced than the effect of lengthening the DNA to 10kbp, suggesting significant restructuring of the internal condensate network by H3K9me3 resulting in part, from its ability to interact with both HP1α and DNA components.

**Figure 4.**
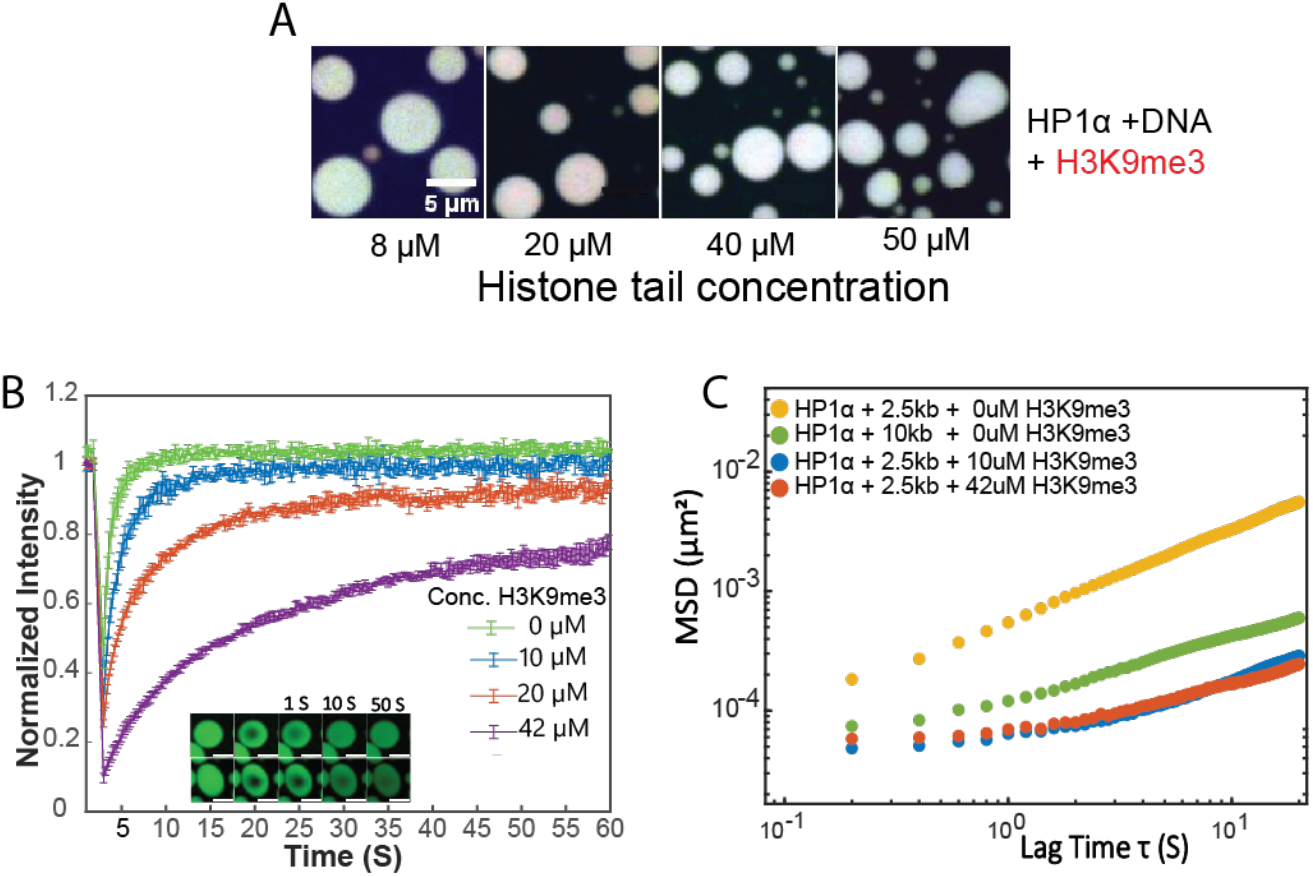
The concentration of H3K9me3 histone peptide fine tunes the material properties of HP1α condensates with DNA. **A)**. Confocal fluorescence images of assemblies of. H3K9me3-Dylight594 peptides (RED) with 20μM HP1α-alexa488(GREEN) and 40nM 2.5kbp DNA-POPO1(BLUE) across a range of histone peptide concentrations. All three components colocalize within condensates containing H3K9me3 peptides. Buffer–70mM KCl, 20mM HEPES, 1mM DTT, pH 7.4 Scale = 5μm. **B)**. Fluorescence recovery over time of Alexa88-HP1α (in 2.5kbp DNA condensates) with increasing H3K9me3 tail concentrations. Scale = 5μm. **C)** MSDs for HP1α-2.5kbp DNA (Yellow) with 10μM H3K9me3 (Blue) and 42μM H3K9me3(Red). MSDs of HP1α-10kbp condensates (Green) shown for comparison.

From a biological perspective, the stability of the morphology of HP1α condensates at different concentrations of the H3K9me3 peptide provides a potential explanation for the fact that heterochromatic states persist in the cells exposed to uniaxial strain. At the same time, the identification of concentration-dependent changes in the internal dynamics and network properties of the HP1α condensates, underscore the enhanced molecular dynamics elicited by altering the relative HP1α:H3K9me3 ratio in response to mechanical forces.

## Discussion

Despite a wealth of knowledge around the hierarchical organization of the genome – from a nucleosome to chromosomes, the mechanisms of genome segregation and the biophysical properties of chromatin sub compartments remain far from understood. Biomolecular phase separation is being increasingly recognized as an underlying mechanism contributing to genome compartmentalization and regulation. Several recent studies have highlighted the ability of nuclear components – including the heterochromatic protein HP1α (13, 14, 17), and histone proteins (16, 21); to form phase separated biomolecular condensates (1, 14, 33). It has thus been proposed that the repressive H3K9me3 epigenetic mark, which is specifically recognized by HP1α, serves as a recruiting element that aids in concentrating and facilitating the phase separation of HP1α. However, whether epigenetic marks, such as H3K9me3 directly contribute to the material state of chromatin has not been explored. It has been shown that HP1α binding to nucleosomes, can further restructure the nucleosome core – which in turn further enhances phase separation (19). Here, to circumvent the complexity of these multifaceted interactions, we developed a reductionist model that allowed for the direct examination of the contribution of the H3K9me3 mark to HP1α phase separation. Using only the histone tails allowed for the decoupling of tail interactions from the contribution of the nucleosome core.

We first established the rheological signatures of the HP1a-DNA condensates in the absence of peptide. Our FRAP results indicating rapid recovery of HP1a, and slower, yet still relatively rapid recovery of DNA across DNA lengths is consistent with previous work (17). Our current work expands our understanding of the material properties of HP1α-DNA condensates, by directly measuring their rheological signatures. Together these FRAP and rheology measurements, which support relatively rapid dynamics and significant elasticity respectively, support the need to consider the relative length and time scales of diffusive dynamics within condensates (34).The slow assembly dynamics of HP1α condensates formed with 10kb long DNA segments, which undergo an apparent transition in morphology, further underscores the viscoelastic nature of DNA condensates.

We then sought to determine whether the presence of histone tails containing either repressive or activating histone marks resulted in the formation of a ternary system comprising H3K9me3/HP1a/DNA. While our FCS data shows enhanced binding between H3K9me3 and HP1α as compared to unmodified H3 tail in the absence of DNA we speculate that this binding specificity might be enhanced in the presence of DNA in the case of the ternary system. It has been suggested that the binding specificity of HP1α towards H3K9me3 peptide is promoted by the presence of non-specific DNA interactions (35) favored by the trimethylation of positively charged lysine residues at position 9 of the histone tail. If electrostatic interaction would be the driving force of the ternary system, then one would expect that H3K9me3 and H3K4me3 peptides would have a similar valence, since they are both characterized by tri-methylation of lysine residues at specific positions.

Remarkably, however. only the presence of H3K9me3 peptides was compatible with the formation of histone/HP1/DNA condensates, while unmodified H3 or H3K4me3 tail, abrogated phase separation of HP1α and DNA. This is of high relevance, both from a biophysical and a biological perspective. At a biophysical level this highlights the importance of structural features of the HP1α protein, such as its chromodomain region of recognition of H3K9me3, but not of H3K4me3 (27). At a biological level, this is of high relevance because it allows for the physical segregation of chromatin compartment with distinctive transcriptional outputs, with HP1α/DNA condensates only associated with transcriptionally repressed heterochromatin. It would be interesting, in future studies to define the potential existence of ternary systems associated with transcriptional activation.

At a cellular level, besides its role in transcriptional repression, heterochromatin also provides a very important role in terms of protection of genomic information from mechanical nuclear strains. During the differentiation of mammalian cells, the deposition of heterochromatic regions at the nuclear periphery determines changes in its mechanical properties (36), with nuclei becoming progressively stiffer over time. When cells are challenged by mechanical forces, they tend to transmit the strain to the nuclear envelope, and pose the threat of nuclear rupture, unless compensatory mechanisms can be enacted. One of these mechanism is the decrease of H3K9me3 from the nuclear periphery, which we detect in microglial cells, consistent with previous results in epithelial cells (24). While protective for the genome, this situation poses an interesting scenario for the HP1α / DNA condensates, which are now faced with lower concentrations of H3K9me3.

In the absence of cellular complexity, the subtle physicochemical differences in the H3 histone peptides, result in significant change in outputs on the material scale in our reductionist system. We show here that even small alterations of the ratio between H3K9me3 and HP1α lead to important changes in the viscoelastic properties of the condensates, which show greater mobility at lower concentration of the repressive mark.

## Concluding remarks

This work highlights the role of HP1α-H3K9me3 interactions as yet another crucial facet of the assembly and functional output of phase-separated heterochromatic states, shedding light on the intricate regulation of constitutive heterochromatin and its response to mechanical cues. Taken together, our results implicate a direct and expanded role for specific histone post-translational modifications in the regulation of material properties of phase separated chromatin bodies.

## Author contributions

PD, SEG designed research and analyzed data related to the biophysics experiment and PD performed the experiments. AVC performed supporting experiments and data analysis. EP designed experiments related to cell work the experiments. PD prepared all the figures and wrote the manuscript, together with PC and SEG.

## Acknowledgements

This work was supported by grants n. NIH R00 NS096217 and AOFSR GRANT12936019 to SEG and grant n. R35NS111604 to PC.

