## Supplementary material for "Modified histone peptides uniquely tune the material properties of HP1α condensates": SI

**Supplemental Document**

I. **Supplemental Figures and legends**
II. **Supplemental Movies and legends**

### I. Supplementary Figures

**SI-1.**

**Figure S1 (A)**. FRAP curves indicating the internal dynamics of 20μM HP1α (1% HP1α-alexa488) in condensates with 2.5kbp(40nM), 4kbp(20nM) and 10kbp (9nM) DNA. **(B)**. FRAP curves indicating the internal dynamics of DNA obtained with YOYO-3 labeled 2.5kbp(40nM), 4kbp(20nM) and 10kbp (9nM) DNA in condensates with 20μM HP1α.

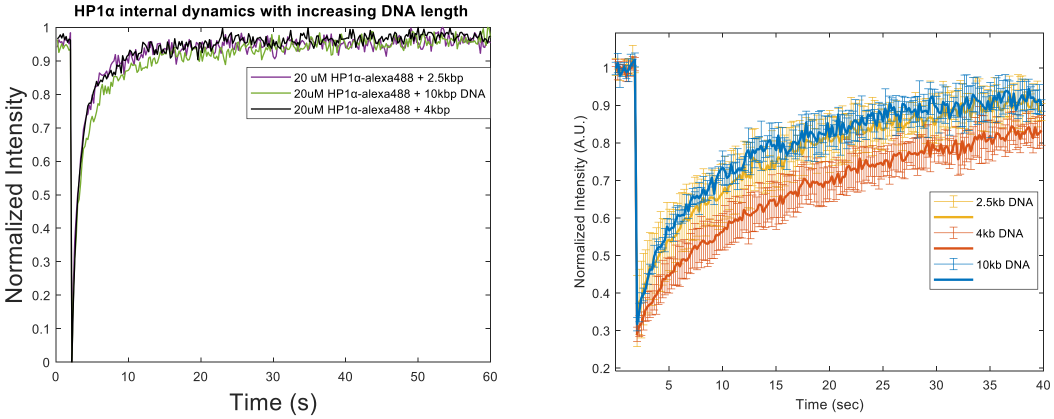


**Figure S1 (A)**. FRAP curves indicating the internal dynamics of 20μM HP1α (1% HP1α-alexa488) in condensates with 2.5kbp(40nM), 4kbp(20nM) and 10kbp (9nM) DNA. **(B)**. FRAP curves indicating the internal dynamics of DNA obtained with YOYO-3 labeled 2.5kbp(40nM), 4kbp(20nM) and 10kbp (9nM) DNA in condensates with 20μM HP1α.

**SI-2 : QC of HP1α and binding specificity towards H3K9me3 (1-22) peptide**

A) SDS PAGE coomasie-stained gel of used HP1α construct (Theoretical molecular weight = 22.6kDA)
B) Molecular weights of unmodified H3 (2353.4 Da), H3K9me3 (2396.3 Da) and H3K4me3 (2395.5 Da) confirmed using MADI.
C) binding affinities of the unmodified (H3) and modified (H3K9me3) peptides quantified using Fluorescence Correlation Spectroscopy. See methods for detailed protocol.

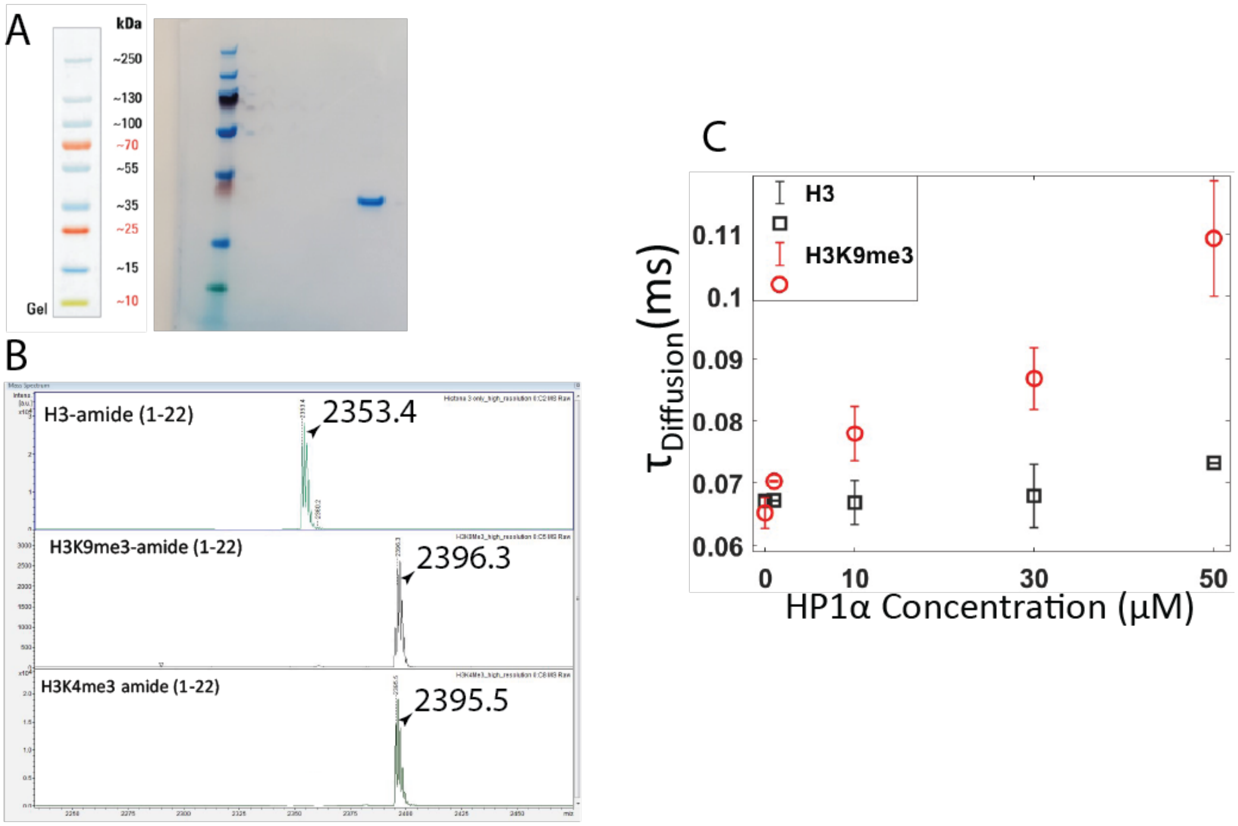

**SI3:.** FRAP curves for aggregate-like assemblies obtained in the case of 20μM HP1α with 42μM Histone H3 (Top, panel A) and H3K4me3 (Bottom, panel B) peptide and YOYO-1 labeled DNA (40nM 2.5kbp) assemblies. HP1α is not localized within the obtained assemblies therefore YOYO-1 (488nm) labeled 2.5kbp DNA is photobleached to estimate the internal dynamics.

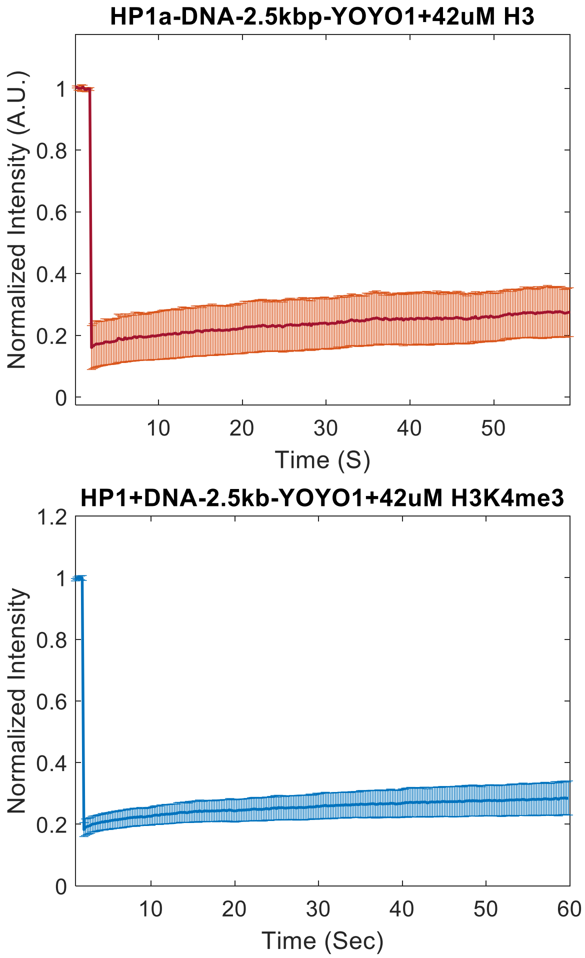

**SI4 : Morphology of assemblies obtained with HP1α, DNA and H3 peptides in distinct combinations.**

A) DIC images of 20 µM HP1α + 42µM histone peptides in the absence of DNA showing no condensate/aggregate formation. Scale = 5µm
B) DIC images of aggregated assemblies obtained with histone peptides in the presence of DNA (2.5kbp represented here) . Scale = 5µm
C) Confocal fluorescence images of assemblies of Histone H3-Dylight594 (RED) with 20µM HP1α-alexa488 (GREEN) and 40nM 2.5kbp DNA-POPO1(BLUE) across a range of histone peptide concentrations. Histone H3 peptide forms assemblies with DNA devoid of HP1α thereby leading to magenta-colored assemblies. Buffer – 70mM KCl, 20mM HEPES, 1mM DTT, pH 7.4 Scale = 5µm

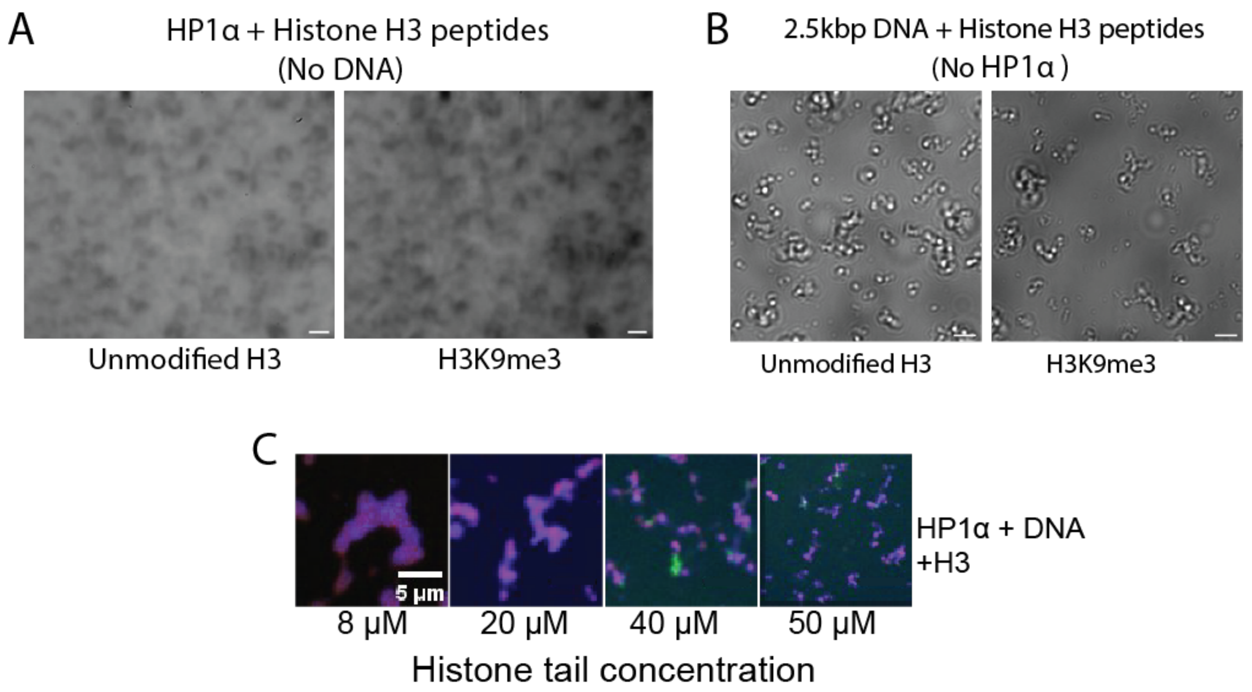

### II. Supplemental Movies

**Movie files can be found here:**

**https://www.dropbox.com/scl/fo/17ymf71h4eqwngv82fhi9/h?rlkey=s0swhszsb5ffptrtaidlgljxm&dl=0**

#### Movie Legends

**Movie S1 (separate file).** Time-dependent transition of 20 μM HP1α + 9 nM 10 kbp DNA asymmetric assemblies into droplet-like morphology over 50 minutes (1 frame = 5min) represented as 1 frame per secondScale bar = 20μM

**Movie S2 (separate file).** Phase-Separated Coacervates composed of 20 μM HP1α, 42 μM H3K9me3 Peptide, and 40 nM 2.5 kbp DNA in 70mM KCl, 20mM HEPES, 1mM DTT, pH 7.4 assembling over 20 mins (1 frame = 2 min) movie speed 1 frame/SScale bar = 20 um.

**Movie S3 (separate file).** Video demonstrating sustained condensate formation with ongoing droplet fusion upon addition of 42 μM H3K9me3 tail from the top (t = 10 seconds) to preformed 20 μM HP1α-alexa 488 + 40 nM 2.5 kbp DNA condensates over a period of 2 min. Movie speed 50 frames/S

**Movie S4** : Video illustrating the liquid-to-aggregate transition of condensates comprising 20 μM HP1α and 40 nM DNA upon addition of 42 μM unmodified H3 peptide from the top (peptide added at t = 24 seconds) over a period of 5 min. Movie speed 50 frames/S

**Movie S5 (Separate file).** FRAP of aggregate-Like assemblies formed with 20 μM HP1α, 40 nM 2.5 kbp-YOYO1 DNA, and 42 μM unmodified H3 peptide demonstrating limited fluorescence recovery after photobleaching.
